# Background color matching influences sexual behavior, growth, and mortality rate in an African cichlid fish

**DOI:** 10.1101/2025.02.05.636662

**Authors:** Travis I. Moore, William G. Bright, William E. Bell, Tessa K. Solomon-Lane, Sebastian G. Alvarado, Peter D. Dijkstra

## Abstract

Phenotypic plasticity allows organisms to adapt to changing environments within their lifetimes. The cost of plastic adaptations may constrain the persistence of plasticity over evolutionary time. One potential cost is the possibility that phenotypic adjustment to specific environments can cause correlated responses that are not necessarily adaptive. Males in the African cichlid *Astatotilapia burtoni* are blue or yellow, and males are able adjust their body coloration to the color of the background, presumably to increase crypsis. To test whether background color influences fitness-related traits, we raised mix-sex groups of juvenile *A. burtoni* to adulthood in yellow or blue tanks. We found that fish in blue tanks were darker and more bluish, whereas fish reared in yellow tanks were paler and more yellow in body coloration. Males, but not females, from blue tanks showed earlier sexual maturation than those held in yellow tanks. However, across the duration of the experiment, there was a higher frequency of females mouthbrooding in groups housed in yellow tanks than those that were housed in blue tanks. In addition, fish in blue tanks exhibited reduced growth rate but higher survivorship relative to their yellow-reared counterparts. Our data suggests that background color affects important fitness-related traits in a color polymorphic cichlid, which may influence the evolution of phenotypic plasticity.

## Introduction

Many organisms can generate adaptive phenotypes in response to changes in environmental conditions in an individual’s lifetime (Agrawal, 2001; Charmantier *et al*., 2008). This ability, called phenotypic plasticity, requires an organism to obtain reliable information about the state of the environment and to translate this information to adaptive changes (Pigliucci *et al*., 2006; Murren *et al*., 2015; Alvarado, 2020). One of the most dramatic examples of phenotypic plasticity is background color matching which has been documented in a broad range of taxa, including insects (Edelaar *et al*., 2017), amphibians (Kindermann *et al*., 2014), reptiles (Corl *et al*., 2018), and fishes (Whiteley *et al*., 2009). Background color matching can occur via changes in the structure and organization of pigmentary cells in the skin to resemble the visual environment and increase crypsis (Sugimoto, 2002; Umbers *et al*., 2014; Ligon & Mccartney, 2016). Cryptic color change is common in many prey species and has been studied extensively (Chiao *et al*., 2011; Smith *et al*., 2016; Liedtke *et al*., 2023).

While plasticity in body coloration may be advantageous in heterogeneous environments (i.e., to increase crypsis across seasons), dramatic phenotypic adjustments can be costly, too (Fischer *et al*., 2014; Murren *et al*., 2015). Describing the nature and the magnitude of these costs can help elucidate under what conditions phenotypic plasticity can evolve and persist. One potential cost of phenotypic plasticity might be caused by correlations between phenotypic plasticity in a trait and other fitness-related traits because of pleiotropy, linkage, or epistasis of genes relevant for the plastic trait (Van Kleunen & Fischer, 2005; Murren *et al*., 2015). More specifically, it is possible that phenotypic adjustments in body coloration can lead to correlated changes that are not necessarily adaptive (Zhang *et al*., 2010; Dijkstra *et al*., 2017; Polo-Cavia & Gomez-Mestre, 2017).

Color change in animals is regulated by a wide range of neuroendocrine systems that often also influence other fitness-related traits. In fish, exposure to a dark background promotes the release of α-melanocyte-stimulating hormone (α-MSH), a melanocortin hormone that increases black body coloration by inducing pigment dispersion and promoting proliferation of melanophores (van Eys & Peters, 1981; Zhang *et al*., 2010). However, α-MSH can also activate melanocortin receptors in the brain and other tissues, thereby affecting a range of physiological and behavioral functions, including metabolism, the stress response, growth rate, and social behavior (Schiöth *et al*., 2005). Other candidate systems for changes in body coloration include melanopsins in pigment containing cells in the integument (Provencio *et al*., 1998). These opsins have been implicated as light sensitive photopigments outside of the retina which can induce color changes in many fishes and frogs but can also lead to behavioral effects such as increased alertness and heightened photosensitivity (Bellingham *et al*., 2002; Bertolesi & McFarlane, 2018). Studies aimed at establishing optimal growth conditions in fish reported that tank color affects a range of traits such as metabolism, behavior, and stress response, which is consistent with the notion that pigmentary adaptation to background color can have correlated effects on other traits [reviewed in (McLean, 2021)]. However, few studies have examined the effect of background color or background color matching on fitness-related traits in the context of phenotypic plasticity (Liedtke *et al*., 2023; Radovanović *et al*., 2023).

In the East African cichlid fish *Astatotilapia burtoni*, males switch between blue and yellow coloration both in the field [Lake Tanganyika, (Fernald & Hirata, 1979)] and the lab (Korzan *et al*., 2008; Dijkstra *et al*., 2017). Coloration is important for mate choice in *A. burtoni* (Dijkstra *et al*., 2024), but conspicuous color morphs run an increased risk of predation (Whitaker *et al*., 2021). Previous work on *A. burtoni* suggests that the reversible coloration allows males to adapt to variable visual environments to reduce the risk of predation, with blue males better adapted to murky algae-rich environments and yellow males better adapted to brighter coastal environments (Fernald & Hirata, 1977; Fang *et al*., 2022). Data from the field suggests that ambient light conditions may vary seasonally in Lake Tanganyika, for example due to upwellings and subsequent algal blooms in certain parts of the lake (Plisnier *et al*., 2023).

Such environmental heterogeneity could favor plasticity in body coloration to maximize cryptic coloration depending on the prevailing environmental conditions. Experimental work in the laboratory showed that *A. burtoni* males can reversibly change color according to the background color with males becoming blue in a blue environment and yellow in a yellow environment (Fang *et al*., 2022). Although color change by exposing individuals to new spectral environments occurs in this species, the potential associated costs with background color adaptation are unknown. In the current study, we assessed how specific background colors affect fitness-related traits in *A. burtoni*. We were specifically interested in how background color influences growth and sexual behaviors during late development as fish are undergoing sexual maturation. To this end, we raised groups of juveniles into adulthood in yellow or blue tanks and quantified color adaptation, growth rate, social behavior, markers of sexual maturation (display of nuptial coloration; mouthbrooding), and survivorship.

## Materials and Methods

### Animals

The cichlid fish *A. burtoni* used in this study were bred from a laboratory population originally derived from Lake Tanganyika (Fernald & Hirata, 1977). Fish were housed in aquaria kept at 28°C with a 12-h light/dark cycle and 10 min each dusk and dawn period to mimic natural settings. Before the experiments, fish were reared in clear 110-liter tanks containing gravel substrate. Continuous water flow and central mechanical and biological filtration occurred throughout the entirety of the experiment. Fish were fed fine granular pellets (Kyorin Food IND. LTD.) and flakes cichlid flakes (Omega Sea LLC). All procedures were approved by Central Michigan University Institutional Animal Care and Use Committee (IACUC approval number 18-10 and 2021-460). A total of 360 fish were used in this study (mean initial length 13.18 mm ± 2.76 mm).

### Experimental housing conditions

A visual of our experimental design is shown in **Figure 1**. Experimental juveniles were collected from rearing tanks (118 liters, 76cm L x 51cm W x 30 cm H, n=7), each containing 40-60 juveniles (∼ age range 2 - 3 months) from 2-4 broods. Juveniles from each rearing tank were randomly split into two groups and assigned to either a tank with blue or yellow background, resulting in a paired experimental tank design (each color treatment, n = 7 tanks). Each experimental tank contained 20 - 28 fish.

**Figure 1.**
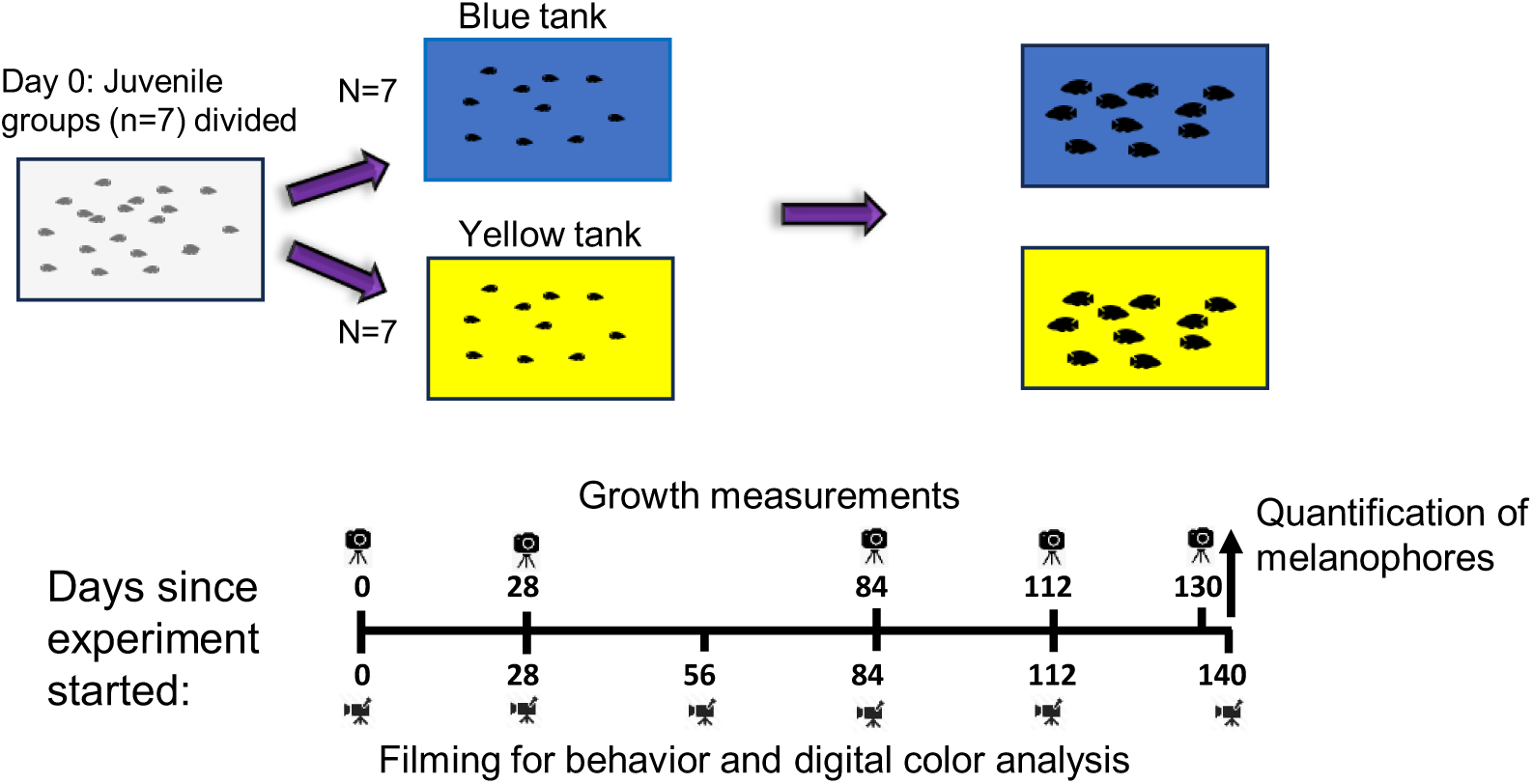
General experiment design. Juvenile *A. burtoni* were raised in mixed-sex groups of 20- 30 individuals for 140 days in yellow or blue tanks. Groups were established when juveniles were 2-3 months of age. In addition to photographing for growth measurements and filming for behavior and digital color analysis, we also recorded the color phenotype of mature males, the number of mouthbrooding females, and mortality rate in each group.

Blue and yellow tanks were identical in dimensions and volume as rearing tanks and were created by fitting blue or yellow acrylic plates to the bottom of tanks and the left, right, and back sides of the tank (U.S. Plastic Corporation, #2050 Blue, #2037 Yellow, 3.175mm thick). Initially, three terracotta flowerpot shards were placed in each experimental tank to provide shelter at the onset of the experiment but were removed after 14 days to reduce early territoriality and aggression in juvenile groups. Yellow or blue colored gravel (Estes’ colored gravel) was placed along the bottoms of corresponding tanks to provide an enriching environment for juveniles. Fish were fed a standardized amount of crushed cichlid flakes (Omega) adjusted to the size and number of fish. We ensured that groups within pairs were fed the exact same amount of food. Juvenile groups were housed in these conditions for 140 days, during which we measured growth, survival, social behavior, the onset of sexual maturation, and body coloration as described below.

### Social behavior and sexual maturation

To evaluate how color treatment influences color development, sexual maturation and the emergence and expression of reproductive behavior, we conducted weekly observations of each tank. During these observations, we quantified the number of males exhibiting bright nuptial coloration as an indicator of sexual maturation in males. For these males, we assigned color phenotype as either blue or yellow. In addition to nuptial coloration, sexually mature males also display egg dummies on the anal fin, and defended a distinct area in their tank (Fraley & Fernald, 1982). We also recorded when spawning began in each group by recording the number of mouthbrooding females.

To quantify social behavior, we filmed each tank every 28 days for a total of six filming sessions with a Cannon EOS 70D. For each recording, the entire tank was filmed for 5-min following a 2-min acclimation period once equipment was set up. All recordings from the group stage were uploaded to BORIS to quantify behaviors (Friard & Gamba, 2016). From behavioral recordings we quantified aggressive behaviors (chases, lateral displays, and border disputes) and courtship behaviors. Aggressive and courtship behaviors were previously described (Fialkowski *et al*., 2021). We scored the total number of occurrences for each behavior in each group because it was not possible to track individual fish in this group setting.

### Survivorship

We recorded the number of fish in each experimental tank on a weekly basis. If a yellow-blue tank pair had differing number of individuals, random individuals would be removed and euthanized from the tank with the higher density to match the density of the tank that experienced any mortality in that week. We performed these corrections since stocking density of tanks can influence growth.

### Growth

The size of all fish was measured five times for each group at days 0, 28, 84, 112, and 130 of the experiment. To reduce the possibility of damage or stress associated with measuring body size, we transferred each group into a small clear plastic container to obtain photographs for digital measurements of body length. The container was placed on a Kaiser RS-1 copy stand measuring 45cm x 50cm with a printed grid of 1cm^2^ squares within the total area. We took photographs of each group using a Pentax K-3 II SLR digital camera attached to a mounting point on the stand fixed 55cm above the fish. Photos were then uploaded to ImageJ to measure the of the body length based on the dorsal view of each juvenile fish following a similar procedure as described previously (Rizzo *et al*., 2017).

### Color quantification

Since stress can induce color change (Nilsson Sköld *et al*., 2013) we did not remove fish from the tanks but instead fish were pulled for digital image analysis from video recordings. At the end of each filming session (every 28 days), we took close-up footage of individual fish within each group for approximately 1 minute for later digital color quantification. We used the standard settings with the autofocus off to prevent the camera from changing focus throughout the video. The fish were filmed at ∼1.0 m away from the aquarium to obtain high resolution pictures with a Kodak Color Separation Guide and Gray Scale (Cat 1527654) in view. In Adobe Photoshop v21.2 we standardized color based on the Kodak Color Separation Guide and Gray Scale using the curves layering technique. This method standardizes collected images and eliminated differences in the lighting between collected images (Bleier *et al*., 2011). We selected images of the fish closest to the front of the aquarium where the lateral of the fish could be clearly seen. In Photoshop, images were converted to CIE L*a*b* after color card correction as described previously for our species (Dijkstra *et al*., 2017). An added advantage of CIE L*a*b* is that it has a single value (b*) that constitutes the balance between blue and yellow, which is of interest to our study species. The images were cut out in Photoshop to remove the background, eyes, and fins. The removed areas were replaced with a uniform, pre-established chroma key of “true green” (CIE L*a*b of 88, −78, 80). The resulting edited images were placed though a JavaScript program that collects the L, a, and b values per individual pixel.

At the end of the 140-day experiment, we took pictures of three to four randomly selected males and females from each group using a stereomicroscope (Leica DM2000LED). Pictures were taken of the caudal fin and the head region. Pictures were analyzed in ImageJ to quantify the melanophores density in the caudal fin, forehead, and mouth regions (upper and lower lips) as described elsewhere (Border *et al*., 2018).

### Data analysis

All analyses were conducted in R v. v3.6.3 (RCore, 2016). We ran linear mixed models (LMMs) or generalized linear mixed models for count data (GLMMs) with tank color and time as fixed or continuous variables and (yellow-blue) pair identity and tank identify as random variables.

To analyze digital color values, we used LMMs to compare b* values between yellow and blue tanks for each observation. For analyzing the presence or absence of territorial males or brooding females over time we used a zero-inflation model (GLMM hurdle models) to examine if territorial males or brooding females appeared sooner in yellow or blue tanks. We compared the number of sexually mature males or brooding females between yellow and blue tanks over time using GLMM with a negative binomial or Poisson distribution (chosen based on AIC values and model diagnostics).

Group-level body size was compared between treatments with LMMs. In these models, we used body size measurements of individual fish and included tank as random effect (we did not track the individual identity of each fish during the experiment). Specific growth rate was analyzed based on group-level body size differences between each growth interval. For the survival analysis, we used a Wilcoxon signed-rank (WSR) test to compare the total number of fish that died between paired yellow and blue tanks. Social behavior was compared between yellow and blue tanks using GLMM with either a negative binomial or Poisson distribution. A significance level (α) of 0.05 was used for all tests. We report mean ± SE for our model estimates.

## Results

### Effect of tank color on body coloration

The male phenotype, in general, matched the background of their tank **(Figure 2a)**, and the proportion of yellow males was significantly higher in groups held in yellow tanks that those held in blue tanks (GLMM, color treatment: 3.50 ± 0.27, z = 12.90, p < 0.00001). Tank color had a significant effect on body coloration as indicated by significant differences in L*a*b* values (**Figure 2b**). The body coloration of fish in yellow tanks was significantly lighter than their blue-reared counterparts as indicated by significantly lower lightness (L*) in blue-reared fish compared to yellow-reared fish (LMM, color treatment: −22.10 ± 6.16, t_14_ = 3.589, p *=* 0.003). Fish in yellow tanks had lower red-green a* values (LMM: −9.00 ± 2.75, t_14_ = −3.267, *p =* 0.00561) but higher blue-yellow b* values (LMM: 30.44 ± 7.30, t_14_ = 4.173, p *=* 0.00094) than fish reared in blue tanks. In addition, at the end of the experiment we found that fish in blue tanks had a significantly higher melanophore density compared to those held in yellow tanks (**Figure 2c**, LMM, color treatment: −1.03 ± 0.13, t_131_ = −8.135, p < 0.0000001) after controlling for significant variation across body regions (LMM, F_132_ = 54.40, p < 0.0000001). In sum, these data suggest fish in blue tanks are more pigmented and more bluish in coloration, whereas fish in yellow tanks are paler and more yellowish.

**Figure 2.**
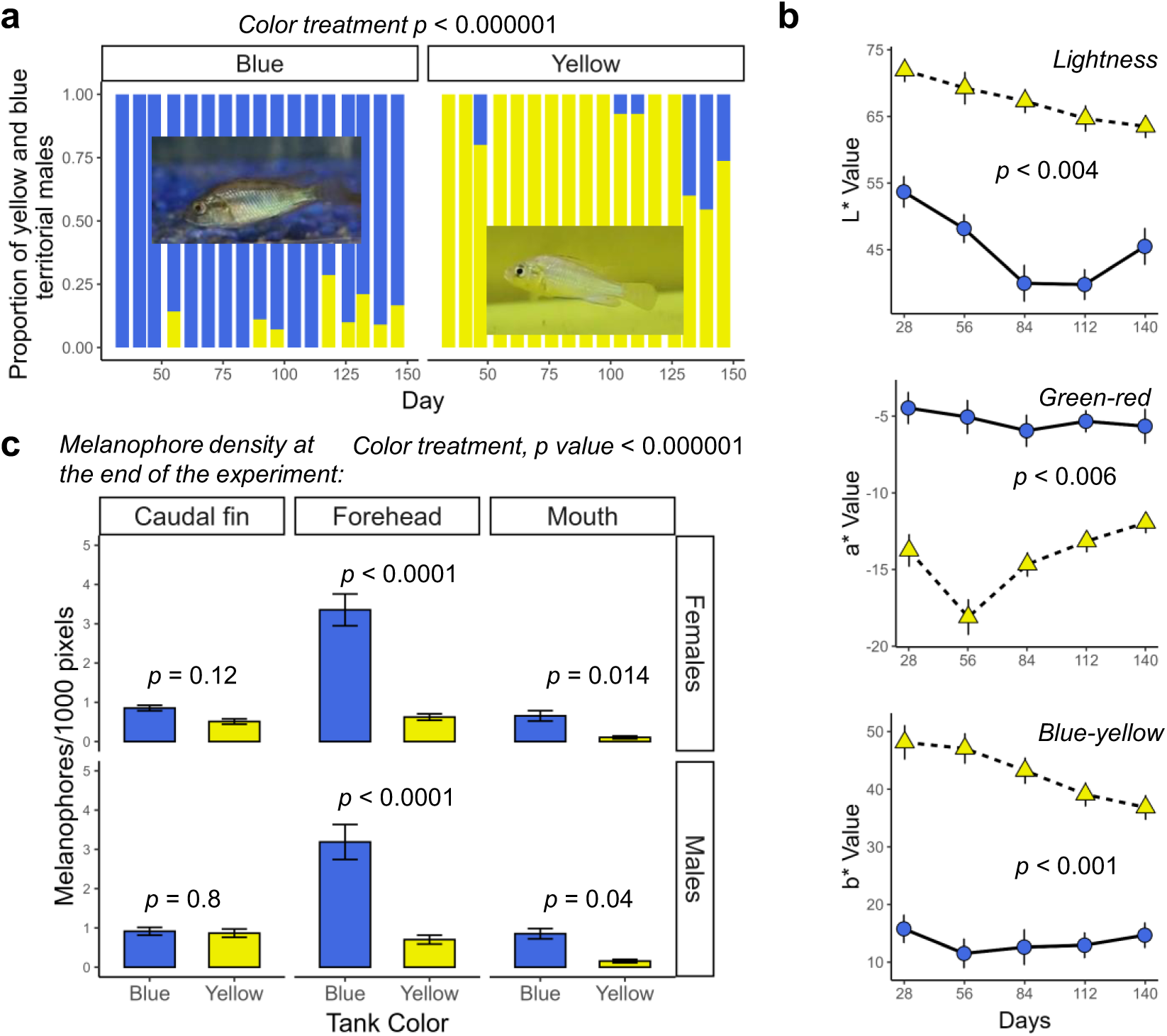
Background color matching. **a.** The proportion of males with blue and yellow nuptial coloration during the experiment. Also shown are representative pictures of males housed in blue or yellow tanks **b**. Melanophore density in different body regions on fish housed in blue or yellow tanks at the end of the experiment. **c**. Digital color quantification of fish yellow and blue tanks. Digital color quantification was performed by capturing images from videos of fish in their tanks and extracting L*a*b* color values. Shown are mean ± se.

### Sexual maturation and behavior

During the first observation (approximately one month after setting up the experiment when fish were ∼ 3-4 months of age), three out of seven blue groups and one out of seven yellow groups had one sexually mature male, as indicated by the expression of nuptial coloration, presence of egg dummies on the anal fin, and increased aggressive behavior. By the end of the experiment, each group had 2 to 4 males expressing nuptial coloration. We found that the first sexually mature male(s) emerged approximately 15 days earlier in groups reared in blue tanks compared to those reared in yellow tanks (**Figure 3a**, LMM, 15.14 ± 5.10, t_7_ = 2.969, p = 0.021). Across the entire experiment, the number of sexually mature males did not differ between yellow and blue tanks after correcting for a significant positive effect of day (**Figure 3b**, GLMM, effect of color treatment: −0.104 ± 0.123, z = −0.844, p = 0.4, effect of day: 0.015 ± 0.002, z = 9.648, p < 0.00001). However, a visual examination of the data (**Figure 3b**) suggests that more sexually mature males were present in blue tanks compared to groups housed in yellow tanks when we began recording sexual behavior (again, observations began approximately one month after establishing groups).

**Figure 3.**
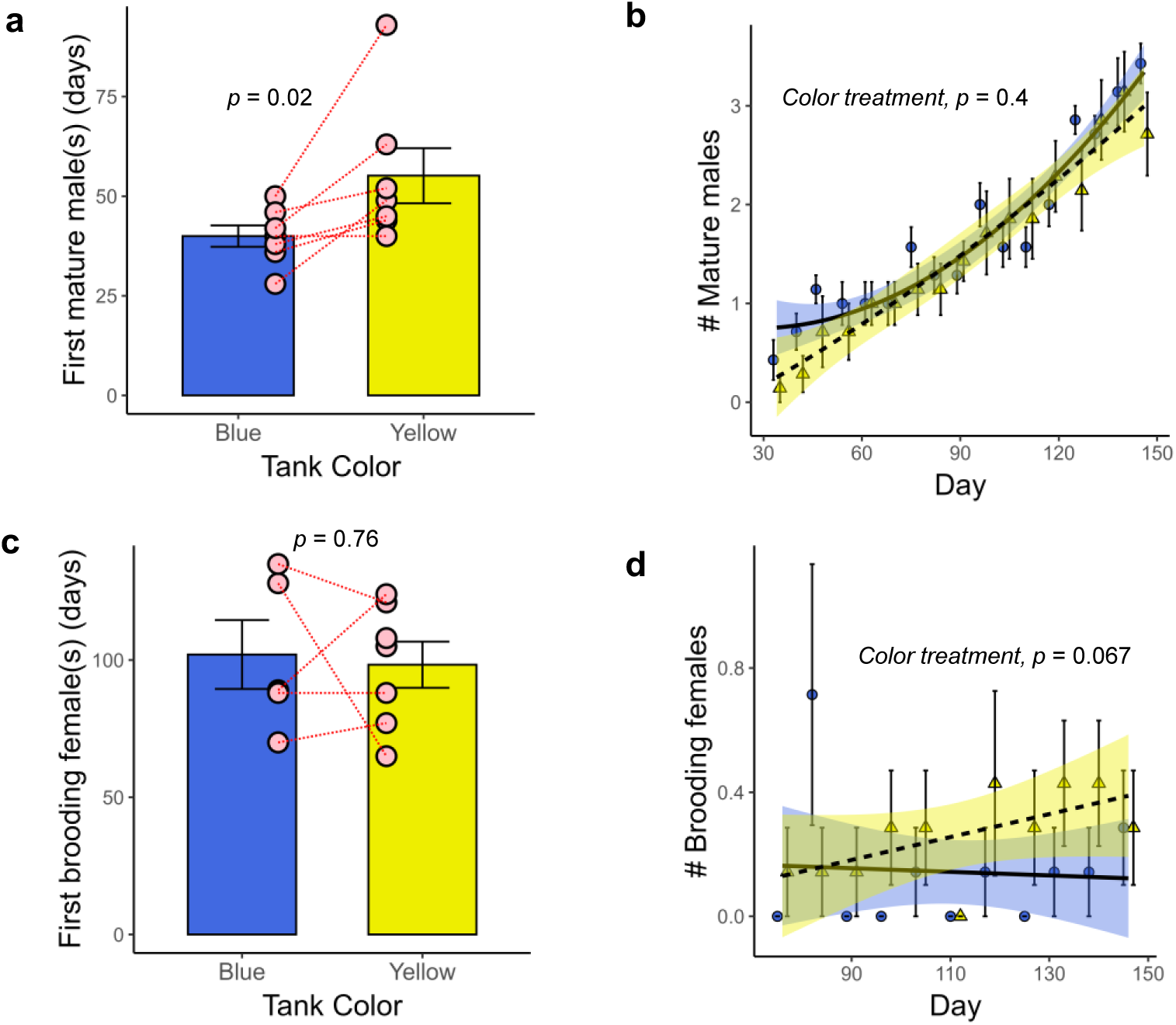
**a.** Sexual maturation as indicated by the onset of males expressing nuptial coloration since the time the experiment was started. **b.** Number of males with nuptial coloration in blue and yellow tanks. **c.** onset of mouthbrooding **d.** Number of brooding females in blue and yellow tanks. In a and c, red lines connect paired groups that originated from the same juvenile tank. Shown are mean ± se.

Mouthbrooding occurred in all groups and started on average 102 days after setting up the experiment (∼5-6 months of age). The onset of mouthbrooding did not vary significantly between yellow and blue-reared groups (**Figure 3c**, LMM, effect of color treatment: −4.08 ± 12.59 t_7_ = −0.324, p = 0.76). However, across the entire experiment, there was a nonsignificant tendency for a higher rate of brooding in females in groups held in yellow tanks compared to those in blue tanks after controlling for the overall increase in breeding over time (**Figure 3d**, GLMM, effect of color treatment: 0.737 ± 0.403, z = 1.830, p = 0.067, effect of day: 0.022 ± 0.007, z = 3.399, p = 0.0007). In line with this, the likelihood of groups containing one or more brooding females was significantly higher in groups reared in yellow tanks compared to those in blue tanks (GLMM zero inflation, color treatment: −0.84 ± 0.43, z = −1.97, p = 0.049).

Tank color did not significantly influence the frequency of chase behaviors (GLMM, color treatment: 0.24791 ± 0.24120, z = 1.028, *p* = 0.304), aggressive display behaviors (GLMM: −0.3456 ± 0.6980, z = −0.495, p = 0.621), or courtship behaviors (GLMM: 0.254 ± 0.449, z = 0.5640, *p* = 0.572) (**Figure 4**). We found that the frequencies of these behaviors changed over time (p values < 0.05), but this effect was not influenced by color treatment (p values > 0.1).

**Figure 4.**
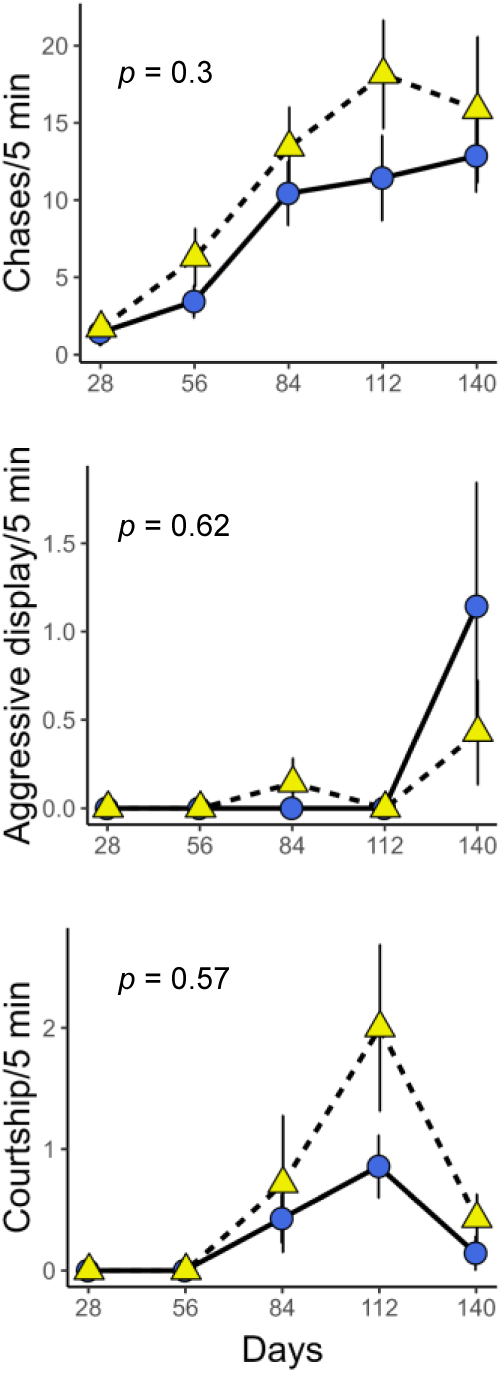
Frequencies of aggression (top), aggressive display (middle), and courtship (bottom) behaviors over time for groups housed in blue and yellow tanks. Shown are mean ± se.

### Growth and survivorship

We found that fish reared in yellow tanks were approximately 4% larger than fish in the blue tanks on the final day when growth was measured (**Figure 5a**, day 130, LMM: 1.20 ± 0.37, t_297_ = 3.265, p = 0.00123). There was no significant size difference between color treatments at the beginning of the experiment (day 0, LMM: −0.36 ± 0.27, t_352_ = −1.372, p = 0.171). When examining specific growth rate in four consecutive intervals during the five-month experiment (calculated based on group averages, **Figure 5b**), we detected significantly higher specific growth rates of yellow-reared fish compared to blue reared fish during the second interval (days 28 through 84 of the experiment, LMM: 0.078 ± 0.023, t_7_ = 3.424, p = 0.011). Specific growth rates did not vary by treatment in the other time intervals (p values > 0.3).

**Figure 5.**
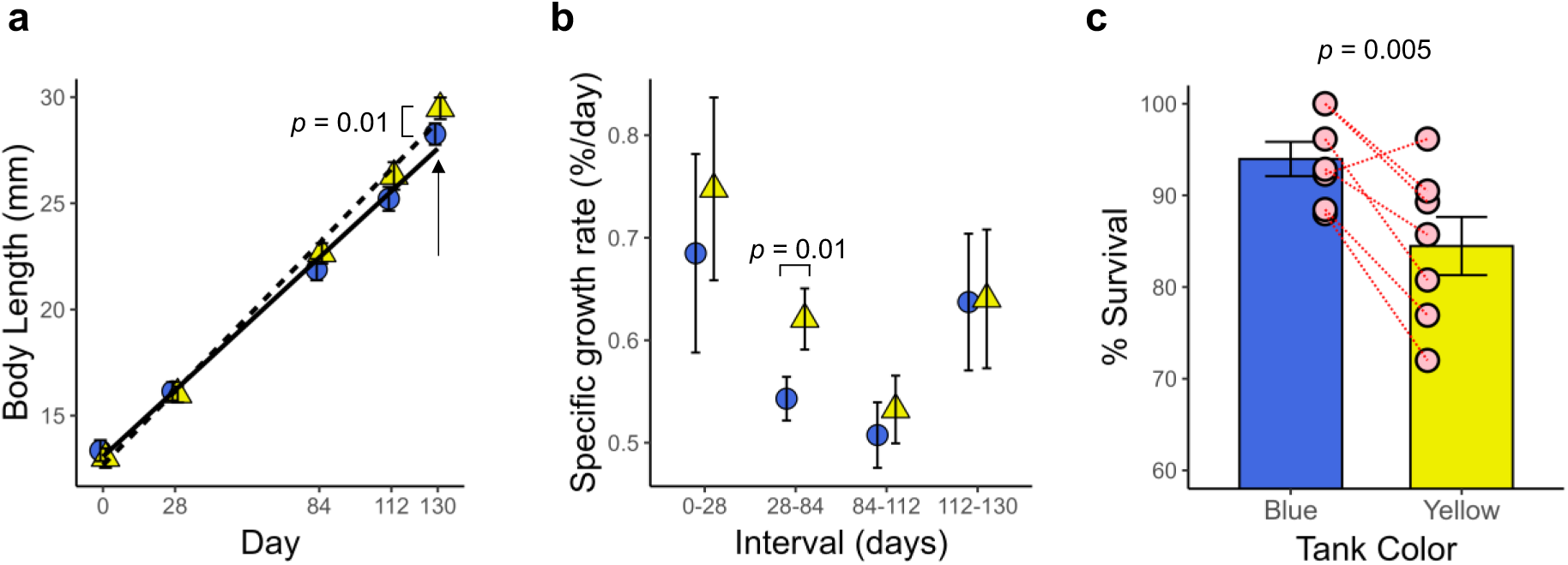
Body size (**a**) and growth rate (**b**) in fish reared in blue and yellow tanks. **b.** Percent survival for fish reared in blue and yellow tanks. Overall percent survival was quantified as the number of surviving fish at the end of the experiment (at day 140) in each tank color divided by the total number of fish in each tank color at the start of the stage. Red lines connect paired groups that originated from the same juvenile tank. Shown are mean ± se.

When considering the cumulative mortality across the experiment, there was significantly more mortality in the yellow tank setting than the blue tank setting (LMM 0.095 ± 0.023, t_7_ = 4.073, p = 0.005) (**Figure 5c**). Groups housed in blue tanks had a mean survivorship of 93.97 ± 1.87% whereas groups in yellow tanks showed a mean survivorship of 84.47 ± 3.17%. Since mortality has been linked to aggression levels in cichlid fish, we also tested whether the overall rate of aggression was linked to mortality rate. Consistent with this idea, there was a significant positive effect of the rate of chases on cumulative mortality after correcting for the effect of color treatment (data not shown: LMM, effect of chases/attacks: 0.015 ± 0.006, t_9_ = 2.492, p = 0.034).

## Discussion

The occurrence of background color adaptation has been widely documented but studies on the effect of specific background colors on important fitness-related traits are relative scarce. In this study, we investigated whether background color during (late) development influences fitness-related traits in the color polymorphic cichlid *A. burtoni* where males are either yellow or blue. As expected, we found that the background color of the tank had a robust influence on body coloration. We then found that fish held in blue tanks exhibited earlier sexual maturation than fish in yellow tanks relative to the emergence of males displaying nuptial (breeding) coloration. Females reared in yellow tanks were breeding more than their counterparts in blue tanks. Finally, we found reduced growth but a tendency for increased survival in groups housed in blue tanks compared to those housed in yellow tanks.

We found that background color affected coloration with fish becoming bluer and more pigmented in blue tanks and fish becoming paler and more yellowish in yellow tanks. We found a similar pattern in the sexually mature males that display bright yellow or blue coloration. Although color matching was not perfect (e.g., blue males did develop in yellow tanks), there were more yellow males in yellow tanks and more blue males in blue tanks. Our findings confirm previous findings showing that male *A. burtoni* adapt their coloration to match their environment (Fang *et al*., 2022). More generally, our findings are consistent with the notion that cichlids, as many other fish species, adjust body pigmentation to the background color, presumably to increase crypsis across heterogeneous environments (Sowersby *et al*., 2015; Takahashi, 2019).

We observed that sexually mature males emerged sooner in groups housed in blue tanks than those housed in yellow tanks. These males are characterized by bright nuptial coloration and often engage in active territorial defense. The expression of nuptial coloration is linked with upregulation of the hypothalamic-pituitary-gonadal axis in haplochromine cichlids (Fraley & Fernald, 1982; Dijkstra *et al*., 2007), suggesting that blue tank color induced more rapid sexual maturation in males compared to those that were housed in yellow tanks. We also note that blue tanks induced increased pigmentation compared to yellow tanks, which is relevant to the expression of nuptial coloration given that is requires hyperpigmentation in several body regions (Dijkstra *et al*., 2007; Border *et al*., 2018). Our findings are consistent with other studies suggesting that tank colors can influence sexual maturation of fishes (Volpato *et al*., 2004; Amiya *et al*., 2008). The onset of mouthbrooding, which is an indicator of sexual maturation in females, did not vary by tank color treatment. However, we found a significantly higher brooding frequency in groups housed in yellow tanks than those housed in blue tanks, as indicated by a marginally nonsignificant higher number of breeding females in yellow tanks and a significantly higher probability of breeding occurring in yellow tanks throughout the entire duration of the experiment. Since egg maturation is metabolically costly (Sawecki *et al*., 2019), we hypothesize the higher growth rate in yellow tanks drives or permits faster egg maturation, allowing females in yellow environments to reproduce more relative to those reared in blue tanks.

Fish in yellow tanks grew faster than fish in blue tanks during all examined intervals, although it was only significant during day 28 through 84 of the experiment. These growth rate differences may have important evolutionary consequences since larger body size confers an advantage in agonistic interactions (Dijkstra *et al*., 2005; Alward *et al*., 2021; Solomon-Lane *et al*., 2022) and mate choice (Thünken *et al*., 2012). The reduced growth rate in blue tanks is consistent with previous work showing that tank color impacts growth (Imanpoor & Abdollahi, 2011; Montajami *et al*., 2012). Given that we observed fainter body coloration in juveniles and young adults housed in yellow tanks, one possibility is that these fish were able to allocate more resources towards growth because they invested less energy towards expressing pigmentation, which is a metabolically expensive process (Polo-Cavia & Gomez-Mestre, 2017; Alfakih *et al*., 2022). However, we found that the cumulative mortality across the duration of the experiment was significantly higher in fish housed in yellow tanks compared to those housed in yellow tanks. The reduced survival rate in yellow tanks is inconsistent with potential metabolic benefits of lower pigmentation in yellow tanks. One possibility is that aggression in yellow tanks led to a higher mortality rate, as has been suggested in other cichlid species (Mohadzir *et al*., 2022). However, there was no significantly higher level of aggression in groups housed in yellow tanks. In addition, although average aggression rates across the entire duration of the experiment were positively correlated with the observed cumulative mortality rates, this effect was the same for blue and yellow tanks. The visual environment can influence the level of chronic stress in different fish species (Kim *et al*., 2016; Costa *et al*., 2017), and it is possible that blue and yellow tanks result in differences in chronic stress, which could impact both growth and survival. Future studies should examine physiological markers of stress, such as cortisol levels in *A. burtoni* reared in yellow tanks compared to those reared in blue tanks.

Cichlid fish rely heavily on visual signals in both intra- and intersexual communication (Seehausen & Van Alphen, 1998; Dijkstra *et al*., 2005; John *et al*., 2021), and variation in the visual environment has been suggested to drive variation in body coloration and even speciation (Seehausen *et al*., 2008; Wright *et al*., 2020). In *A. burtoni*, we previously reported that blue males are sexually more attractive, while yellow males are superior fighters when competing for mating territories (Dijkstra *et al*., 2024). It will be interesting to see how background color adaptation and sexual selection interact and influence color expression and plasticity in this species. For example, it is possible that plasticity driven by the environment may conflict with the optimal body coloration in the context of mate choice or male-male competition [(Kelley *et al*., 2016), see also (Smith *et al*., 2016)]. Integrating these fitness costs in combination with background color specific effects on fitness-related traits reported in this study may contribute to our understanding of how the physical environment combined with sexual selection influence the evolution of phenotypic plasticity in body coloration. We note that the reported effects of background color on fitness-related traits could be an intrinsic cost of plasticity if these effects are detrimental to Darwinian fitness. However, in the current study we did not test if background color adaptation was linked to actual fitness costs, which would be difficult to measure. It is also possible that the observed differential background color effects on fitness-related traits are adaptive. For example, life history theory predicts that earlier sexual maturation is typically favored when resources are unpredictable (Stearns, 1992). In Lake Tanganyika, bluer or murkier visual environments occur when lake productivity and the risk of harmful algal blooms creating anoxic conditions are high (Plisnier *et al*., 2023). It is possible that under these circumstances (high food availability but also high unpredictability relative to risk of dying), it may be advantageous to undergo faster sexual maturation. Finally, in the current study, we did not study how color change, that is background color adaptation after transferring males to another tank color, influences fitness-related traits. Such an approach would specifically test the cost associated with color change (Radovanović *et al*., 2023), and this will be a focus in future studies. Finally, early life experiences can influence behavior later in life (Solomon-Lane & Hofmann, 2019), and we are currently examining how the visual environment influences male behavior and phenotype in adult life.

In summary, background color affected a variety of fitness-related traits in *A. burtoni*, including sexual maturation, breeding frequency, growth, and survivorship. Our results suggest that distinct visual environments may differentially impact important fitness-related traits, which could play a role in the evolution of color plasticity. In general, the adaptive significance of plasticity in body coloration according to background color depends on the relative cost and benefits on specific backgrounds as well as the spatial and temporal heterogeneity of the visual environment (Wente & Phillips, 2003; Corl *et al*., 2018; Fang *et al*., 2022). Given the importance of body coloration in social communication, it will be interesting to see how background color adaptation and social selection interact to influence the evolution of phenotypic plasticity in body coloration. Finally, phenotypic plasticity is thought to play an important role in evolutionary diversification [e.g., plasticity may precede or facilitate adaptive evolution (Levis & Pfennig, 2016)], and will be interesting to see how plasticity in body coloration contributes to adaptive diversification in nuptial coloration and other traits in the East African cichlid radiations (Malinsky *et al*., 2015; Kratochwil *et al*., 2018; Wright *et al*., 2020).

## Acknowledgements

We thank members of the Dijkstra lab for providing helpful comments to earlier drafts of the manuscript. This research was supported by grants from Central Michigan University (Undergraduate Summer Scholars Awards to WGB and WB; a Graduate Student Award to TIM; and a Faculty Research and Creative Endeavors Award to PDD), a grant from the Grants for Laboratory Animal Science (GLAS) program of the American Association for Laboratory Animal Science (AALAS), and a grant from the National Institute of General Medical Sciences (NIGMS, R15GM150286).

## References

Agrawal, A.A. 2001. Ecology: Phenotypic plasticity in the interactions and evolution of species. Science. 294: 321–326.

Alfakih, A., Watt, P.J. & Nadeau, N.J. 2022. The physiological cost of colour change: evidence, implications and mitigations. J. Exp. Biol. 225: jeb210401.

Alvarado, S.G. 2020. Molecular plasticity in animal pigmentation: Emerging processes underlying color changes. Integr. Comp. Biol. 60: 1531–1543.

Alward, B.A., Cathers, P.H., Blakkan, D.M., Hoadley, A.P. & Fernald, R.D. 2021. A behavioral logic underlying aggression in an African cichlid fish. Ethology 127: 572–581.

Amiya, N., Amano, M., Yamanome, T., Yamamori, K. & Takahashi, A. 2008. Effects of background color on GnRH and MCH levels in the barfin flounder brain. Gen. Comp. Endocrinol. 155: 88–93.

Bellingham, J., Whitmore, D., Philp, A.R., Wells, D.J. & Foster, R.G. 2002. Zebrafish melanopsin: Isolation, tissue localisation and phylogenetic position. Mol. Brain Res. 107: 128–136.

Bertolesi, G.E. & McFarlane, S. 2018. Seeing the light to change colour: An evolutionary perspective on the role of melanopsin in neuroendocrine circuits regulating light-mediated skin pigmentation. Pigment Cell Melanoma Res. 31: 354–373.

Bleier, M., Riess, C., Beigpour, S., Eibenberger, E., Angelopoulou, E., Tröger, T., et al. 2011. Color constancy and non-uniform illumination: Can existing algorithms work? In: Proceedings of the IEEE International Conference on Computer Vision.

Border, S.E., Piefke, T.J., Fialkowski, R.J., Tryc, M.R., Funnell, T.R., Deoliveira, G.M., et al. 2018. Color change and pigmentation in a color polymorphic cichlid fish. Hydrobiologia 832: 175–191.

Charmantier, A., McCleery, R.H., Cole, L.R., Perrins, C., Kruuk, L.E.B. & Sheldon, B.C. 2008. Adaptive phenotypic plasticity in response to climate change in a wild bird population. Science. 320: 800–803.

Chiao, C.C., Wickiser, J.K., Allen, J.J., Genter, B. & Hanlon, R.T. 2011. Hyperspectral imaging of cuttlefish camouflage indicates good color match in the eyes of fish predators. Proc. Natl. Acad. Sci. U. S. A. 108: 9148–9153.

Corl, A., Bi, K., Luke, C., Challa, A.S., Stern, A.J., Sinervo, B., et al. 2018. The genetic basis of adaptation following plastic changes in coloration in a novel environment. Curr. Biol. 28: 2970–2977.e7.

Costa, D.C., Mattioli, C.C., Silva, W.S., Takata, R., Leme, F.O.P., Oliveira, A.L., et al. 2017. The effect of environmental colour on the growth, metabolism, physiology and skin pigmentation of the carnivorous freshwater catfish Lophiosilurus alexandri. J. Fish Biol. 90: 922–935.

Dijkstra, P.D., Funnell, T.R., Fialkowski, R.J., Piefke, T.J., Border, S.E., Aufdemberge, P.M., et al. 2024. Sexual selection may support phenotypic plasticity in male coloration of an African cichlid fish. Proc. R. Soc. B Biol. Sci. 291: 20241127. The Royal Society.

Dijkstra, P.D., Hekman, R., Schulz, R.W. & Groothuis, T.G.G. 2007. Social stimulation, nuptial colouration, androgens and immunocompetence in a sexual dimorphic cichlid fish. Behav. Ecol. Sociobiol. 61: 599–609.

Dijkstra, P.D., Maguire, S.M., Harris, R.M., Rodriguez, A.A., DeAngelis, R.S., Flores, S.A., et al. 2017. The melanocortin system regulates body pigmentation and social behaviour in a colour polymorphic cichlid fish. Proc. R. Soc. B Biol. Sci. 284: 20162838.

Dijkstra, P.D., Seehausen, O. & Groothuis, T.G.G. 2005. Direct male-male competition can facilitate invasion of new colour types in Lake Victoria cichlids. Behav. Ecol. Sociobiol. 58: 136–143.

Edelaar, P., Baños-Villalba, A., Escudero, G. & Rodríguez-Bernal, C. 2017. Background colour matching increases with risk of predation in a colour-changing grasshopper. Behav. Ecol. 28: 698–705.

Fang, W., Blakkan, D., Lee, G., Bashier, R., Fernald, R.D. & Alvarado, S.G. 2022. DNA methylation of the endothelin receptor B makes blue fish yellow. bioRxiv, doi: 10.1101/2022.09.27.509821.

Fernald, R.D. & Hirata, N.R. 1977. Field study of Haplochromis burtoni: Quantitative behavioural observations. Anim. Behav. 25: 964–975.

Fernald, R.D. & Hirata, N.R. 1979. The ontogeny of social behavior and body coloration in the african cichlid fish haplochromis burtoni. Zeitschrif für Tierpsychologie 50: 180–187.

Fialkowski, R., Aufdemberge, P., Wright, V. & Dijkstra, P. 2021. Radical change: temporal patterns of oxidative stress during social ascent in a dominance hierarchy. Behav. Ecol. Sociobiol. 75: 43.

Fischer, B., van Doorn, G.S., Dieckmann, U. & Taborsky, B. 2014. The evolution of age-dependent plasticity. Am. Nat. 183: 108–125.

Fraley, N.B. & Fernald, R.D. 1982. Social control of developmental rate in the African cichlid, Haplochromis burtoni. Z. Tierpsychol. 60: 66–82.

Friard, O. & Gamba, M. 2016. BORIS: a free, versatile open-source event-logging software for video/audio coding and live observations. Methods Ecol. Evol. 7: 1325–1330.

Imanpoor, M.R. & Abdollahi, M. 2011. Effects of Tank Color on Growth, Stress Response and Skin Color of Juvenile Caspian Kutum Rtilus frisii Kutum. Glob. Vet. 6: 118–125.

John, L., Rick, I.P., Vitt, S. & Thünken, T. 2021. Body coloration as a dynamic signal during intrasexual communication in a cichlid fish. BMC Zool. 6: 1–13.

Kelley, J.L., Rodgers, G.M. & Morrell, L.J. 2016. Conflict between background matching and social signalling in a colour-changing freshwater fish. R. Soc. Open Sci. 3: 160040.

Kim, B.S., Jung, S.J., Choi, Y.J., Kim, N.N., Choi, C.Y. & Kim, J.W. 2016. Effects of different light wavelengths from LEDs on oxidative stress and apoptosis in olive flounder (Paralichthys olivaceus) at high water temperatures. Fish Shellfish Immunol. 55: 460–468.

Kindermann, C., Narayan, E.J. & Hero, J.M. 2014. The neuro-hormonal control of rapid dynamic skin colour change in an amphibian during amplexus. PLoS One 9: e114120.

Korzan, W.J., Robison, R.R., Zhao, S. & Fernald, R.D. 2008. Color change as a potential behavioral strategy. Horm. Behav. 54: 463–70.

Kratochwil, C.F., Liang, Y., Gerwin, J., Woltering, J.M., Urban, S., Henning, F., et al. 2018. Agouti-related peptide 2 facilitates convergent evolution of stripe patterns across cichlid fish radiations. Science. 362: 457–460.

Levis, N.A. & Pfennig, D.W. 2016. Evaluating “plasticity-first” evolution in nature: key criteria and empirical approaches. Trends Ecol. Evol. 31: 563–574.

Liedtke, H.C., Lopez-Hervas, K., Galván, I., Polo-Cavia, N. & Gomez-Mestre, I. 2023. Background matching through fast and reversible melanin-based pigmentation plasticity in tadpoles comes with morphological and antioxidant changes. Sci. Rep. 13: 12064.

Ligon, R.A. & Mccartney, K.L. 2016. Biochemical regulation of pigment motility in vertebrate chromatophores: A review of physiological color change mechanisms. Curr. Zool. 62: 237–252.

Malinsky, M., Challis, R.J., Tyers, A.M., Schiffels, S., Terai, Y., Ngatunga, B.P., et al. 2015. Genomic islands of speciation separate cichlid ecomorphs in an East African crater lake. Science. 350: 1493–1498.

McLean, E. 2021. Fish tank color: An overview. Aquaculture 530: 735750.

Mohadzir, S., Rahmah, S., Rasdi, N.W., Jalilah, M., Ghaffar, M.A., Chang, Y., et al. 2022. Intraspecific aggression in the jewel cichlid Hemichromis bimaculatus reared under different background colours. Aquac. Res. 53: 6407–6413.

Montajami, S., Nkoubin, H., Mirzaie, F.S. & Sudagar, M. 2012. Influence of Different Artificial Colors of Light on Growth Performance and Survival Rate of Texas Cichlid Larvae (Herichthys cyanoguttatus). World J. Zool. 7: 232–235.

Murren, C.J., Auld, J.R., Callahan, H., Ghalambor, C.K., Handelsman, C.A., Heskel, M.A., et al. 2015. Constraints on the evolution of phenotypic plasticity: Limits and costs of phenotype and plasticity. Heredity. 115: 293–301.

Nilsson Sköld, H., Aspengren, S. & Wallin, M. 2013. Rapid color change in fish and amphibians - function, regulation, and emerging applications. Pigment Cell Melanoma Res. 26: 29–38.

Pigliucci, M., Murren, C.J. & Schlichting, C.D. 2006. Phenotypic plasticity and evolution by genetic assimilation. J. Exp. Biol. 209: 2362–7.

Plisnier, P.D., Cocquyt, C., Cornet, Y., Poncelet, N., Nshombo, M., Ntakimazi, G., et al. 2023. Phytoplankton blooms and fish kills in Lake Tanganyika related to upwelling and the limnological cycle. J. Great Lakes Res. 49: 102247.

Polo-Cavia, N. & Gomez-Mestre, I. 2017. Pigmentation plasticity enhances crypsis in larval newts: Associated metabolic cost and background choice behaviour. Sci. Rep. 7: 39739.

Provencio, I., Jiang, G., De Grip, W.J., Pär Hayes, W. & Rollag, M.D. 1998. Melanopsin: An opsin in melanophores, brain, and eye. Proc. Natl. Acad. Sci. U. S. A. 95: 340–345.

Radovanović, T.B., Petrović, T.G., Gavrilović, B.R., Despotović, S.G., Gavrić, J.P., Kijanović, A., et al. 2023. What coloration brings: Implications of background adaptation to oxidative stress in anurans. Front. Zool. 20: 1–12.

Rizzo, A.A., Welsh, S.A. & Thompson, P.A. 2017. A paired-laser photogrammetric method for in situ length measurement of benthic fishes. North Am. J. Fish. Manag. 37: 16–22.

Sawecki, J., Miros, E., Border, S.E. & Dijkstra, P.D. 2019. Reproduction and maternal care increase oxidative stress in a mouthbrooding cichlid fish. Behav. Ecol. 40: 1662–1671.

Schiöth, H.B., Haitina, T., Ling, M.K., Ringholm, A., Fredriksson, R., Cerdá-Reverter, J.M., et al. 2005. Evolutionary conservation of the structural, pharmacological, and genomic characteristics of the melanocortin receptor subtypes. Peptides 26: 1886–1900.

Seehausen, O., Terai, Y., Magalhaes, I.S., Carleton, K.L., Mrosso, H.D.J., Miyagi, R., et al. 2008. Speciation through sensory drive in cichlid fish. Nature 455: 620–626.

Seehausen, O. & Van Alphen, J.J.M. 1998. The effect of male coloration on female mate choice in closely related Lake Victoria cichlids (Haplochromis nyererei complex). Behav. Ecol. Sociobiol. 42: 1–8.

Smith, K.R., Cadena, V., Endler, J.A., Kearney, M.R., Porter, W.P. & Stuart-Fox, D. 2016. Color change for thermoregulation versus camouflage in free-ranging lizards. Am. Nat. 188: 668– 678.

Solomon-Lane, T.K., Butler, R.M. & Hofmann, H.A. 2022. Vasopressin mediates nonapeptide and glucocorticoid signaling and social dynamics in juvenile dominance hierarchies of a highly social cichlid fish. Horm. Behav. 145: 105238.

Solomon-Lane, T.K. & Hofmann, H.A. 2019. Early-life social environment alters juvenile behavior and neuroendocrine function in a highly social cichlid fish. Horm. Behav. 115: 104552.

Sowersby, W., Lehtonen, T.K. & Wong, B.B.M. 2015. Background matching ability and the maintenance of a colour polymorphism in the red devil cichlid. J. Evol. Biol. 28: 395–402.

Stearns, S. 1992. The evolution of life histories. p. 249. Oxford University Press, Oxford, UK.

Sugimoto, M. 2002. Morphological color changes in fish: Regulation of pigment cell density and morphology. Microsc. Res. Tech. 58: 496–503.

Takahashi, T. 2019. Colour variation of a shell-brooding cichlid fish from Lake Tanganyika. Hydrobiologia 832: 193–200.

Thünken, T., Meuthen, D., Bakker, T.C.M. & Baldauf, S.A. 2012. A sex-specific trade-off between mating preferences for genetic compatibility and body size in a cichlid fish with mutual mate choice. Proc. R. Soc. B Biol. Sci. 279: 2959–2964.

Umbers, K.D.L., Fabricant, S.A., Gawryszewski, F.M., Seago, A.E. & Herberstein, M.E. 2014. Reversible colour change in Arthropoda. Biol. Rev. 89: 820–848.

van Eys, G.J.J.. & Peters, P.T.. 1981. Evidence for a direct role of α-MSH in morphological background adaptation of the skin in Sarotherodon mossambicus. Cell Tissue Res. 217: 361–372.

Van Kleunen, M. & Fischer, M. 2005. Constraints on the evolution of adaptive phenotypic plasticity in plants. New Phytol. 166: 49–60.

Volpato, G.L., Duarte, C.R.A. & Luchiari, A.C. 2004. Environmental color affects Nile tilapia reproduction. Brazilian J. Med. Biol. Res. 37: 479–483.

Wente, W.H. & Phillips, J.B. 2003. Fixed green and brown color morphs and a novel color-changing morph of the Pacific tree frog Hyla regilla. Am. Nat. 162: 461–73.

Whitaker, K.W., Alvarez, M., Preuss, T., Cummings, M.E. & Hofmann, H.A. 2021. Courting danger: socially dominant fish adjust their escape behavior and compensate for increased conspicuousness to avian predators. Hydrobiologia 848: 3667–3681.

Whiteley, A.R., Gende, S.M., Gharrett, A.J. & Tallmon, D.A. 2009. Background matching and color-change plasticity in colonizing freshwater sculpin populations following rapid deglaciation. Evolution. 63: 1519–1529.

Wright, D.S., van Eijk, R., Schuart, L., Seehausen, O., Groothuis, T.G.G. & Maan, M.E. 2020. Testing sensory drive speciation in cichlid fish: Linking light conditions to opsin expression, opsin genotype and female mate preference. J. Evol. Biol. 33: 422–434.

Zhang, C., Song, Y., Thompson, D. a, Madonna, M. a, Millhauser, G.L., Toro, S., et al. 2010. Pineal-specific agouti protein regulates teleost background adaptation. Proc. Natl. Acad. Sci. U. S. A. 107: 20164–71.

